# Bursting emerges from the complementary roles of neurons in a four-cell network

**DOI:** 10.1101/2021.08.05.455351

**Authors:** Akira Sakurai, Paul S. Katz

## Abstract

Reciprocally inhibitory modules that form half-center oscillators require mechanisms for escaping or being released from inhibition. The central pattern generator (CPG) underlying swimming by the nudibranch mollusc, *Dendronotus iris*, is composed of only four neurons that are organized into two competing modules of a half center oscillator. In this system, bursting activity in left-right alternation is not driven by any of the neurons but is an emergent property of the network as a whole. We found that the unique synaptic actions and membrane properties of the two neurons in each module (Si2 and Si3) play complementary roles in generating stable bursting in this network oscillator. Only Si2 evokes fast and strong inhibition of its contralateral counterpart, the termination of which initiates post-inhibitory rebound in the Si3 of that module. Only Si3 is responsible for the rebound excitation because it has a hyperpolarization-activated inward current. Within each module, the synaptic actions and membrane properties of the two neurons complement each other: Si3 excites Si2, which then feeds back slow inhibition to Si3, terminating the burst. Using dynamic clamp, we showed that the magnitude of the slow inhibition sets the period of the oscillator. Thus, the synaptic actions of Si2 provide the hyperpolarization needed for the other module to rebound stably, whereas the membrane properties of Si3 in each module cause it to rebound first and excite Si2 and maintain the burst until the cycle repeats and the other module becomes active.

## Introduction

Since the original proposal of the half-center oscillator (HCO) concept by Brown (1911), reciprocal inhibition between two modules has been considered to be the fundamental architecture of network oscillators (Grillner and El Manira 2015; Kiehn 2016; Moult et al. 2013). However, reciprocal inhibition by itself is not sufficient to maintain stable rhythmic bursting. In many systems, endogenous or conditional bursting properties contribute to maintaining the alternation of half-center oscillators (Arshavsky et al. 1986; Cymbalyuk et al. 2002; Kiehn et al. 1996; Li et al. 2010; Reith and Sillar 1998; Selverston and Miller 1980; Wallen and Grillner 1987). A mechanistic cellular-level understanding of network-driven oscillations in a central pattern generator (CPG) has been difficult to determine because of the large numbers of neurons in most network oscillators. The CPG underlying swimming in the nudibranch mollusc, *Dendronotus iris* is a network oscillator consisting of only four neurons (Sakurai and Katz 2016). Here we examined synaptic actions and membrane properties of these two pairs of bilaterally represented neurons in the generation of stable bursting activity.

*Dendronotus* swims by rhythmically flexing its body in alternation from left to right (Sakurai et al. 2011). The rhythmic motor pattern originates from a swim CPG that consists of neurons swim interneurons numbered Si2 and Si3 (Fig. 1A,B) (Sakurai and Katz 2016). Both Si2 and Si3 reciprocally inhibit their contralateral counterparts contributing to the left-right alternation (Fig. 1B). However, the two modules of the HCO, which we label α and β here, consist of the contralateral Si2 and Si3, such that the left Si2 and the right Si3 fire in phase with each other as module α and the right Si2 fires with the left Si3 as module β (Fig. 1C*i*). A strong excitatory synapse from Si3 to the contralateral Si2 and electrical coupling between the pair mediates their synchronous firing in each module (Fig. 1C*ii*). In this study, we investigated the mechanisms underlying the transition of activity between the two modules of the swim CPG. We found that each of the two neurons in a module contributes differently to burst initiation and termination of the cycle. Specifically, the combined action of a slow synapse and active membrane properties provide dynamic features needed for this half-center oscillator to function.

**Figure 1.**
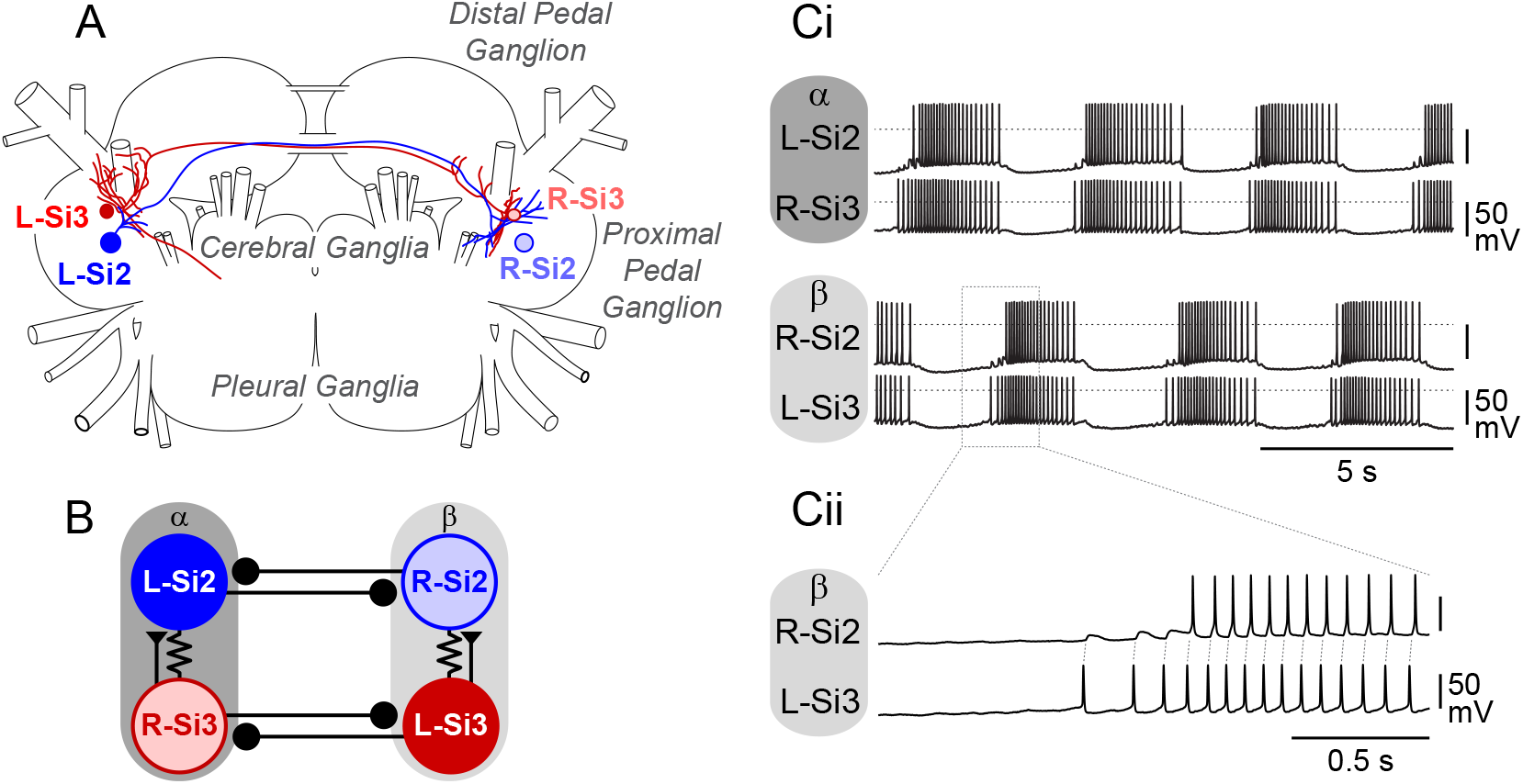
The central pattern generator (CPG) underlying the swimming behavior of *Dendronotus iris*. ***A***. Schematic drawing of the cell body locations and the axonal projections of the swim CPG neurons, Si2 (blue) and Si3 (red), in the *Dendronotus* brain, showing Cerebral, Pleural, and two Pedal (proximal and distal) ganglia based on Sakurai and Katz (2016). Si2 and Si3 have their cell bodies in the proximal pedal ganglion. They each project an axon to the contralateral pedal ganglion through one of the two pedal commissures. Only L-Si2 and R-Si3 axon projections are depicted. ***B***. The previously published connectivity of the *Dendronotus* swim CPG. Each module of the half-center oscillator that constitutes the swimming circuit is composed of Si2 and Si3 with cell bodies on opposite sides. Module α consists of the left Si2 and right Si3, and module β consists of the right Si2 and left Si3. Lines terminating in triangles indicate excitatory synapses, filled circles inhibitory synapses. Resistor symbols indicate electrical connections. ***C***. The swim motor pattern recorded intracellularly from all four swim interneurons (***Ci***). Si2 and Si3 in the same module fire bursts of action potentials together. The two modules exhibit alternating bursts. A portion of the traces indicated by the dotted box is enlarged in ***Cii***. L-Si3 spikes evoke EPSPs in the R-Si2, leading to one-for-one spikes.

## Materials and Methods

### Animal collection, maintenance, and dissection

Specimens of *Dendronotus iris* (Mollusca; Gastropoda; Nudibranchia), 6-20 cm long, were collected on the Pacific coast of North America by Monterey Abalone Company (Monterey, CA) and Living Elements Ltd (Delta, BC, Canada). The animals were kept in artificial seawater tanks at 10-12°C with a 12:12 light/dark cycle.

Before the surgery for brain isolation, the animal was anesthetized by injecting 0.33 M magnesium chloride solution (20-50 mL) into the body cavity, and an incision was made through the body wall near the esophagus. Then all nerve roots exiting the brain were severed, and the brain (Fig. 1A) was isolated by severing the esophagus. The isolated brain was pinned with the dorsal side up on the bottom of a Sylgard-lined dish and perfused at a rate of 0.5 ml/min with normal saline (mM: 420 NaCl, 10 KCl, 10 CaCl_2_, 50 MgC1_2_, 11 D-glucose, 10 HEPES, pH 7.6) or artificial seawater (Instant Ocean, Mentor, OH). The brain was chilled down to 4°C, and the connective tissue and the brain sheath were removed with forceps and fine scissors. After desheathing, the temperature was raised to 10°C for electrophysiological experiments.

### Electrophysiology

Intracellular recordings were performed using 15-35 MΩ glass microelectrodes filled with a solution containing 2 M potassium acetate and 0.2 M potassium. Each electrode was connected to the headstage of Axoclamp 2B amplifier (Molecular Devices, Sunnyvale, CA). The output signals of the amplifier were digitized at a sampling frequency of >2 kHz using a 1401Plus or Micro1401 A/D converter (Cambridge Electronic Design, Cambridge, UK). Data acquisition and analysis were performed with Spike2 software (CED, Cambridge, UK) and SigmaPlot (Jandel Scientific, San Rafael, CA).

To investigate the effects of depolarization or hyperpolarization of the swim interneurons on the swim rhythm, a positive or negative current pulse (from −4 nA to 4 nA) was injected directly into the neurons via bridge-balanced microelectrodes. To investigate monosynaptic connection between swimming interneurons, high-divalent cation (Hi-Di) saline was applied to raise the spike threshold and suppress spontaneous neural firing. The composition of Hi-Di saline (in mM) was 285 NaCl, 10 KCl, 25 CaCl_2_,125 MgCl_2_,11 D-glucose, 10 HEPES, pH 7.6. To measure the reversal potentials of synaptic potential or synapticc current, the membrane potential was manipulated by injecting a steady current through an additional microelectrode placed in the same neuron. In some experiments, calcium-free Hi-Di saline was used to block chemical synaptic transmission. In low calcium Hi-Di saline, CaCl_2_ was replaced by MgCl_2_.

### Dynamic clamp

To artificially enhance or counteract the Si2-evoked slow IPSPs in Si3, we performed dynamic clamping with the software StdpC (Kemenes et al. 2011) as described previously (Sakurai and Katz 2017; 2016; 2019; Sakurai et al. 2014b). Each Si3 was impaled with two microelectrodes, one for membrane potential recording and the other for current injection (see Fig. 10Ai). Every time Si2 generated an action potential that exceeded 0 mV, the current injected into the postsynaptic Si3, *I*_stim_, was calculated using a first order kinetics model of the release of neurotransmitter (Destexhe et al. 1994; Kemenes et al. 2011; Sharp et al. 1996):

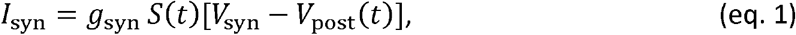

where *S*(*t*) is the instantaneous synaptic activation, *g*_syn_ is the maximum synaptic conductance, *V*_syn_ is the reversal potential (−70 mV) of the synapse. To counteract Si2-evoked synaptic potentials, a negative value was used for *g*_syn_.

**Figure 2.**
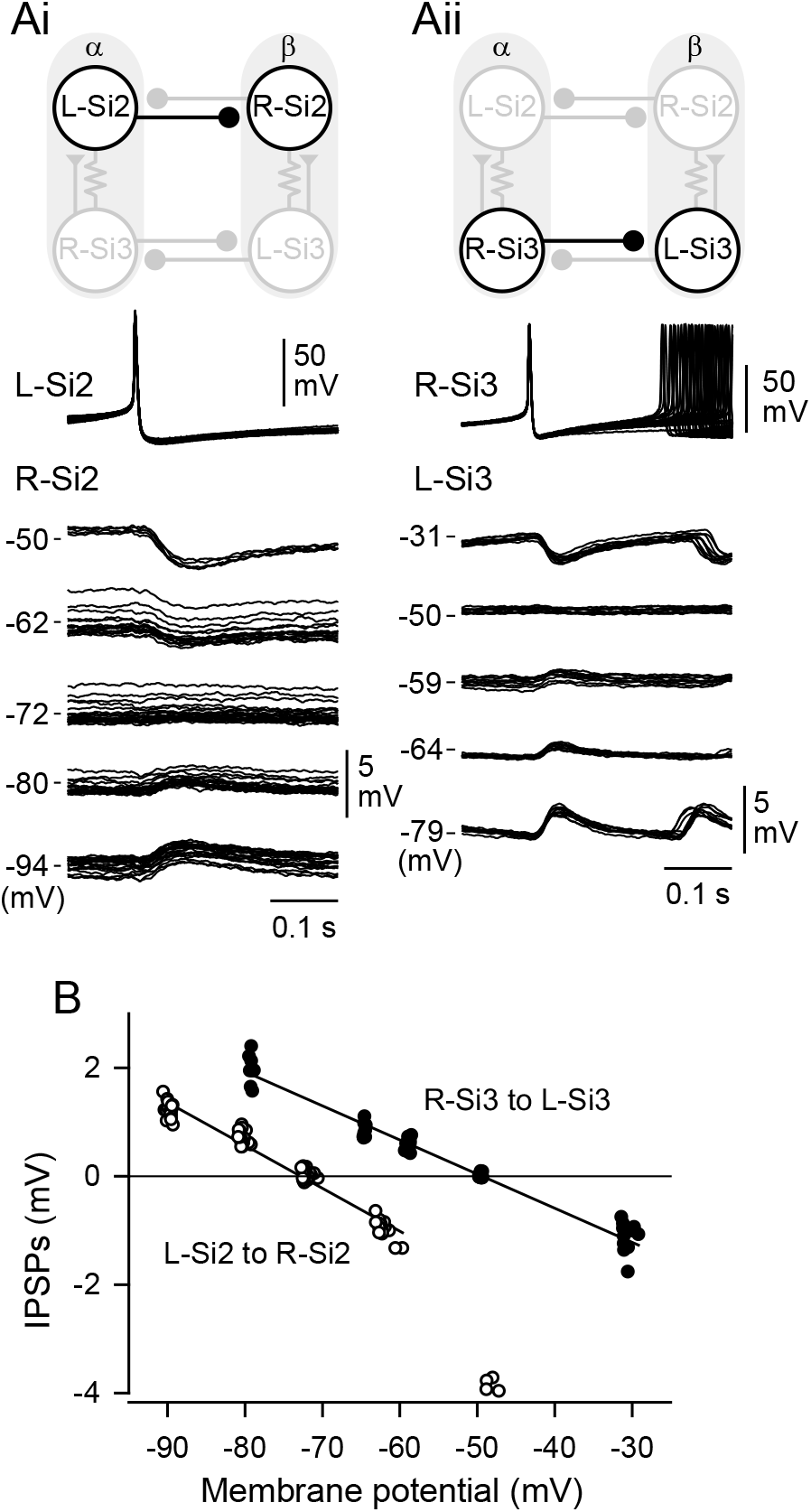
The inhibitory synaptic potentials generated by Si2 and Si3 have different reversal potentials. ***A***. Schematic diagrams of synaptic connections in the swim CPG and overlaid traces of L-Si2 to R-Si2 (*Ai*) and L-Si3 to R-Si3 (*Aii*) synaptic potentials in Hi-Di saline. One electrode was inserted into L-Si2 (*Ai*) or R-Si3 (*Aii*) to record the presynaptic action potentials, while two electrodes (one for voltage recording and the other for current injection) inserted into R-Si2 (*Ai*) or L-Si3 (*Aii*), respectively. Positive or negative current was injected into the postsynaptic cell (R-Si2 or L-Si3) to change its membrane potential. The numbers on the left side of the traces indicate the postsynaptic membrane potential. Ten traces were overlaid for each voltage level. Traces were triggered at the peak of presynaptic action potentials. ***B***. The relationships between the amplitude of the IPSPs and the membrane potential of the postsynaptic cell are shown. The values used are derived from the data shown in *A*. The white circles indicate the Si2-evoked IPSPSs in the contralateral Si2, and the filled circles are the Si3-evoked IPSPs in the contralateral Si3. The regression lines were drawn for the linear portion only.

**Figure 3.**
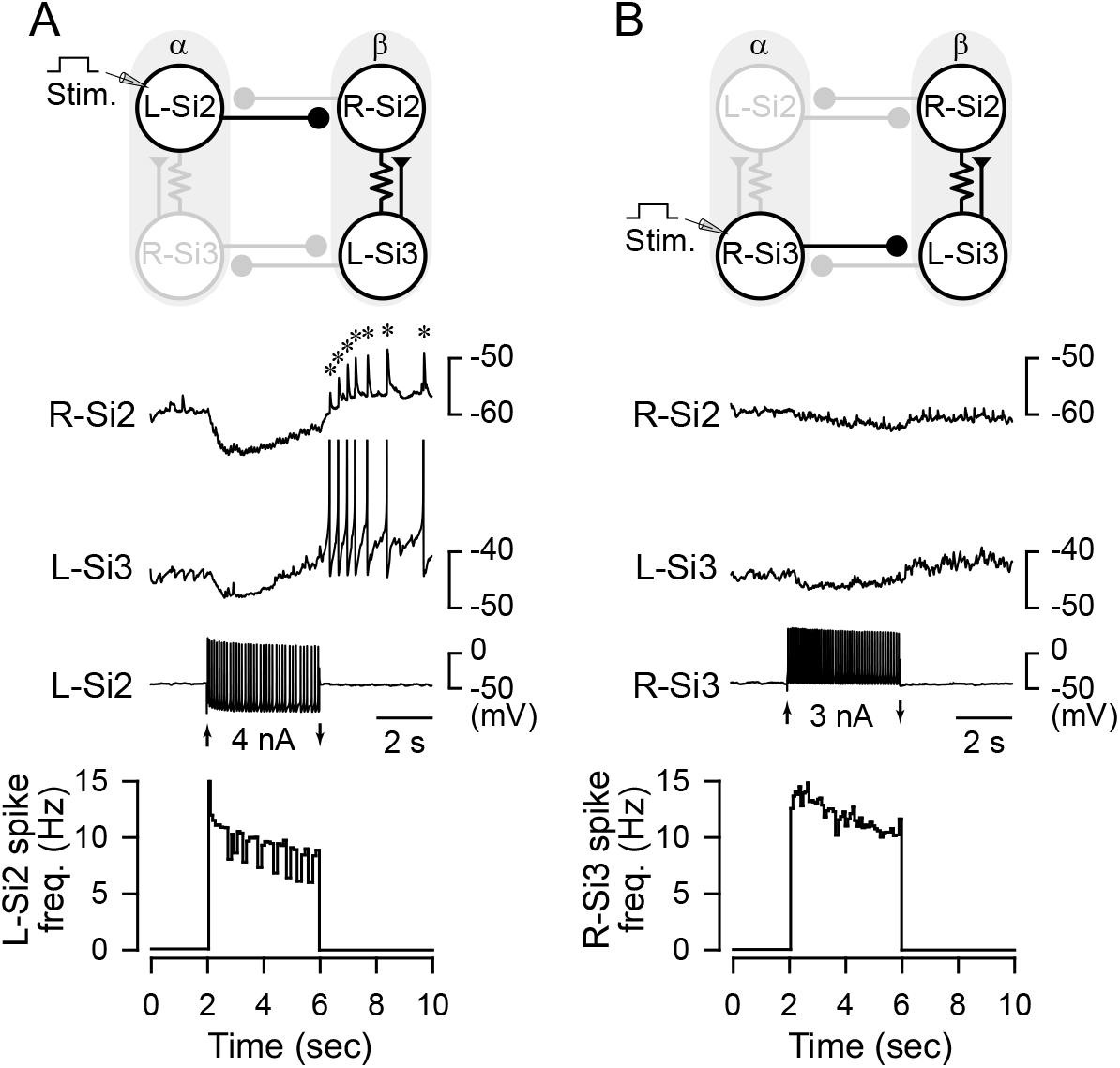
An Si2 spike train causes a post-inhibitory rebound discharge in Si3 of the contralateral module. Top: schematic of circuit with recorded neurons highlighted. Middle: Simultaneous intracellular microelectrode recordings. Bottom: Plot of instantaneous spike frequency of stimulated neuron. ***A***. Depolarization of the L-Si2 with a 4 sec, 4 nA current pulse, caused it to fire spike train with a peak frequency of 15 Hz, that declined to about 10 Hz. Si2 spiking evoked hyperpolarizing responses in Si2 and Si3 of the other module, followed by post-inhibitory discharges in the L-Si3 that evoked EPSPs in the R-Si2 (asterisks). ***B***. Depolarization of the R-Si3 with a 4 sec, 3 nA current pulse caused is to fire with a peak frequency of 14 Hz, which decayed to 10 Hz. Si3 spiking produced only small hyperpolarization in the neurons of the other module and caused no rebound discharge.

**Figure 4.**
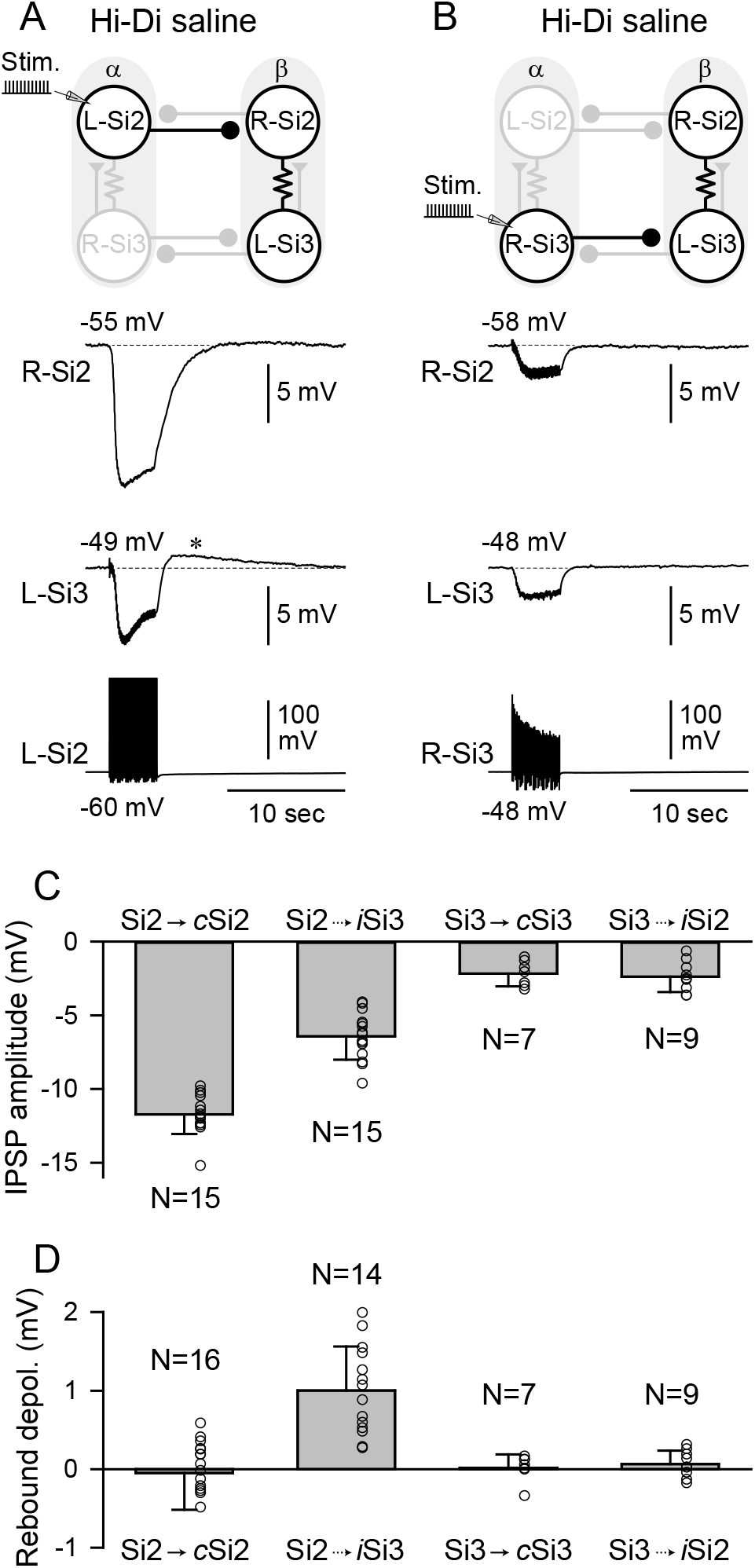
A Si2 spike train causes a sag depolarization and a post-inhibitory rebound depolarization in postsynaptic Si3. ***A***. A 4 sec, 10 Hz spike train evoked with brief current pulses (10 nA, 20 msec) in the L-Si2 caused hyperpolarization of both Si2 and Si3 in Module β. After the stimulation, the membrane potential of the postsynaptic Si2 slowly recovered back to the resting potential, whereas Si3 showed a faster recovery and a rebound depolarization (asterisk), which lasted for 10 to 15 sec. ***B***. A 10-Hz spike train in the R-Si3 caused hyperpolarization of both Si2 and Si3 in Module β. The amplitude of hyperpolarization was smaller than those evoked by Si2 in ***A***, and there was no rebound depolarization after the stimulus. ***C***. Amplitudes of the Si2- and Si3-evoked hyperpolarization in contralateral (c) and ipsilateral (*i*) postsynaptic neurons. Each bar represents mean±SD (N=7-15). ***D***. Amplitudes of the post-inhibitory rebound depolarization. Each bar represents mean±SD (N=7-16).

**Figure 5.**
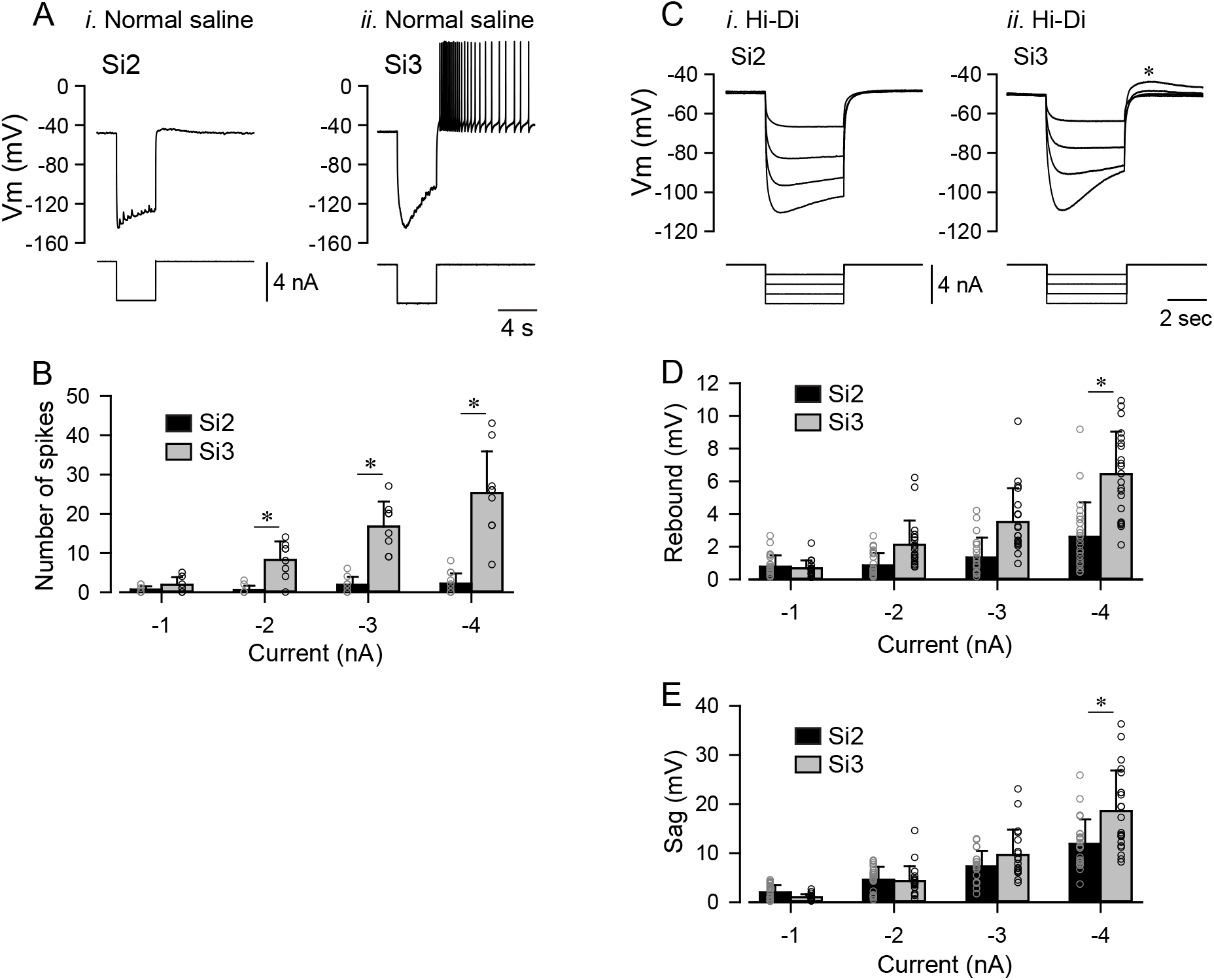
Si3 but not Si2 showed sag potential and rebound depolarization. ***A***. Membrane potential responses (upper traces) of Si2 (*i*) and Si3 (*ii*) to a hyperpolarizing current pulse (−4 nA for 4 sec, lower trace) in normal saline. ***B***. The numbers of spikes in Si2 (black) and Si3 (gray) evoked after hyperpolarizing current pulses of various amplitudes (−1, −2, −3, and −4 nA). Each bar represents mean±SD (Si2, N = 9-12; Si3, N = 7-10). The asterisk indicates significant difference (see text). ***C***. Overlaid membrane potential responses (upper traces) of Si2 (*i*) and Si3 (*ii*) to hyperpolarizing current pulses (4 sec, −1 to −4 nA, lower traces) in Hi-Di saline. The asterisk indicates a depolarizing overshoot. ***D***. Amplitudes of peak rebound depolarization with respect to resting membrane potential in Si2 (black) and Si3 (gray) after hyperpolarizing current injections (−1 to −4 nA, 4 sec). The asterisk indicates significant difference (see text). ***E***. The amplitudes of sag depolarization, measured from peak hyperpolarization to value at end of pulse, in Si2 (black) and Si3 (gray) during the hyperpolarizing current injection (−1 to −4 nA, 4 sec). The asterisk indicates significant difference (see text).

**Figure 6.**
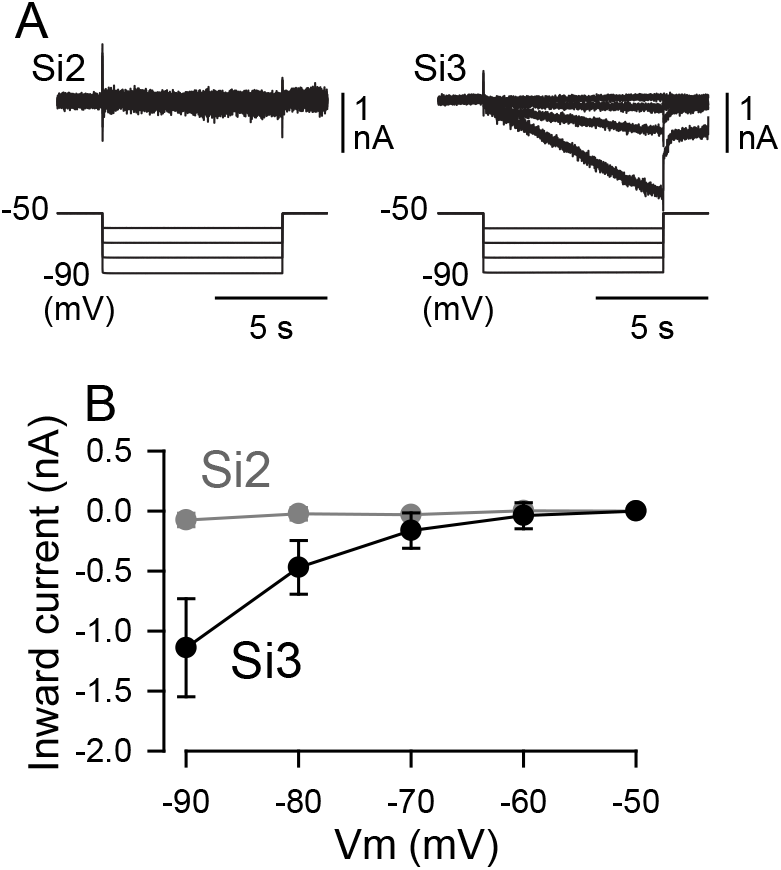
Si3, but not Si2, exhibited a hyperpolarization-activated inward current. ***A***. Membrane current responses (upper traces) and command voltage pulses (bottom traces). The membrane potentials of Si2 (left) and Si3 (right) were held at −50 mV under voltage clamp and then were stepped to −60, −70, −80, −90 mV). Each neuron was impaled with two electrodes (one for voltage recording and the other for current injection). All voltage-clamp experiments were performed in Hi-Di saline. ***B***. The amplitudes (nA, mean±SD; Si2, N=5; Si3, N = 8) of membrane current are plotted against the command voltage values (mV).

**Figure 7.**
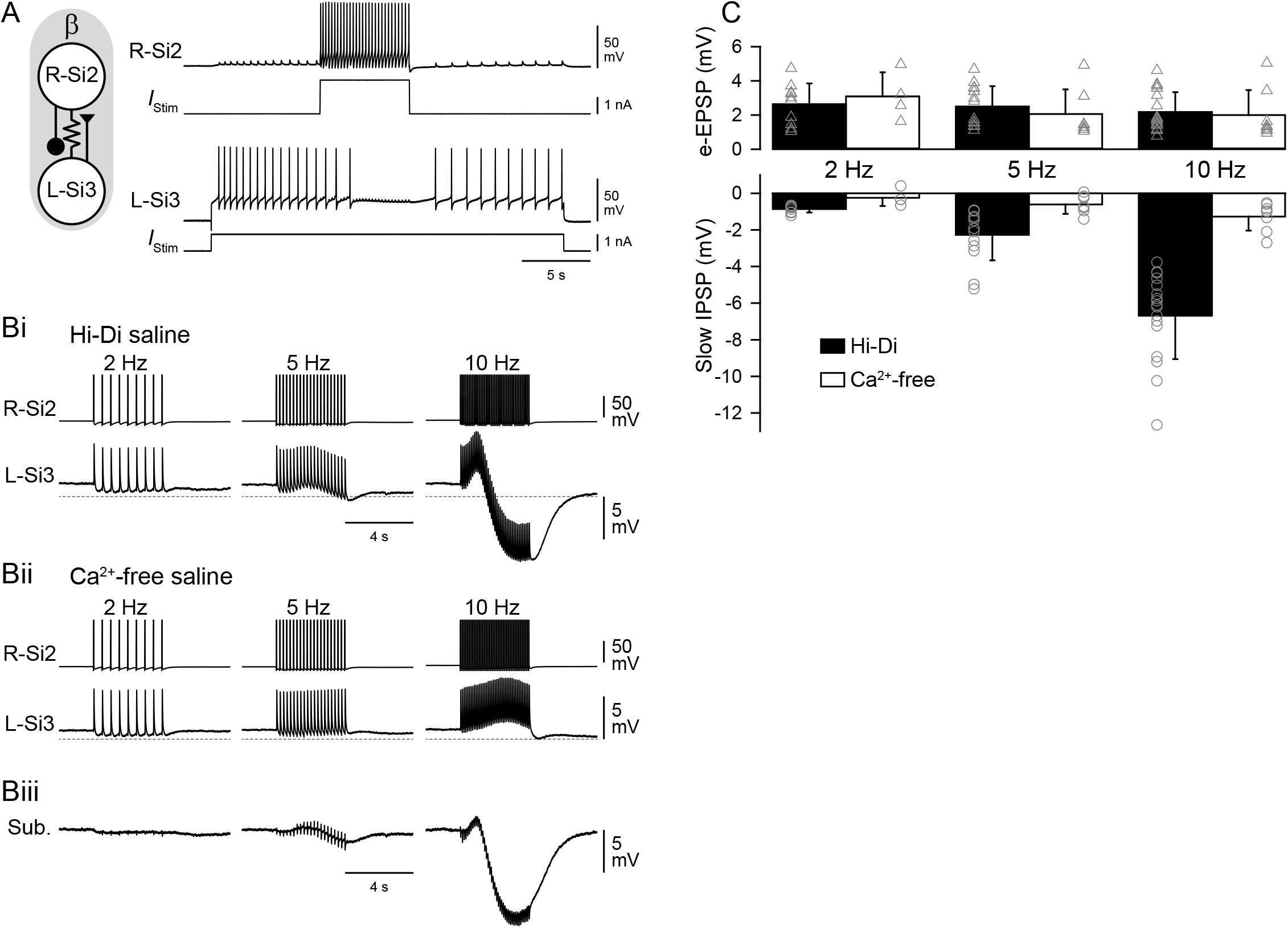
A spike train in Si2 evokes a slow inhibitory synaptic potential in Si3 in the same module. ***A***. An Si2 spike train inhibited ongoing spiking activity of Si3. Simultaneous intracellular records of Si2 and Si3 and the injected current traces. Tonic spiking in L-Si3 was evoked by 25 sec, 1 nA current pulse. A 5 sec, 2nA current pulse in the R-Si2 inhibited spiking of the L-Si3. The schematic on the left shows synaptic interactions between Si2 and Si3 within the module. ***B***. A spike train of Si2 (upper trace) produced complex membrane potential responses of Si3 (lower trace) in the same module, part of which was mediated by chemical transmitter release. Trains of action potentials in R-Si2 were evoked by injecting repetitive current pulses (7-15 nA, 20 msec) at 2, 5, and 10 Hz. ***Bi***. In Hi-Di saline, the R-Si2 spikes produced sharp electrotonic EPSPs in L-Si3, which overrode on a slow hyperpolarizing potential. ***Bii***. Ca^2+^-free saline blocked the slow hyperpolarizing potential, leaving the electrotonic EPSPs relatively unchanged. ***Biii***. Subtraction of the Bii waveform from the Bi waveform revealed the slow hyperpolarizing component that was largely blocked by the Ca^2+^-free saline. ***C***. Amplitudes of electrotonic EPSPs (upper graph) and the slow hyperpolarizing potentials (slow IPSPs; lower graph) recorded in Hi-Di saline (black bars) and in Ca^2+^-free Hi-Di saline (white bars). Ca2+-free saline diminished the slow IPSPs but had little effect on the electrotonic EPSPs. Bars are mean±SD.

**Figure 8.**
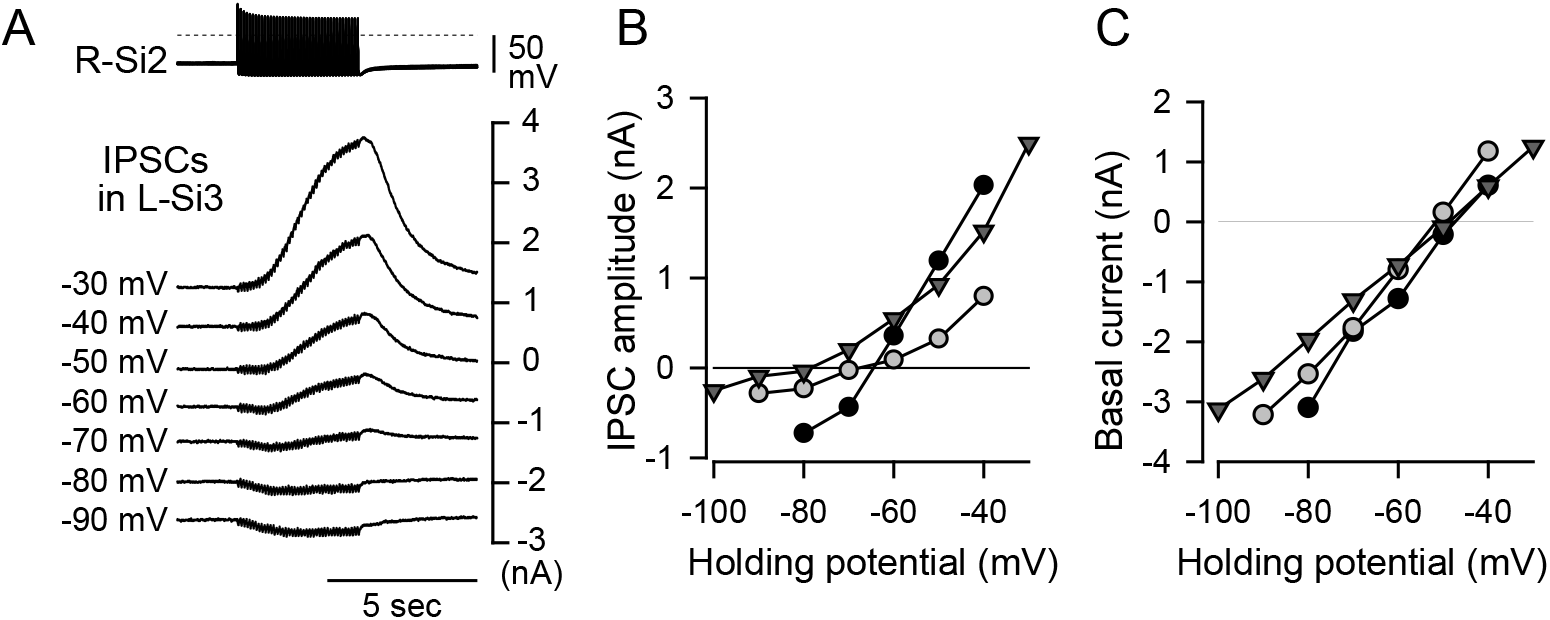
R-Si2 evoked a slow outwardly rectifying current in L-Si3. ***A***. Example traces of the synaptic currents at different holding potentials. Spike trains were evoked in R-Si2 by injecting repetitive current pulses (10 nA, 20 msec) at 10 Hz. Seven traces of the Si2 spike trains are overlaid and shown at the top, whereas the slow IPSPs in L-Si3, which was voltage-clamped at various potential levels, are shown below. The holding potentials (from −90 mV to −30 mV) are shown on the left side of the trace. ***B***. The amplitudes of the slow IPSPs plotted against the holding potential of the postsynaptic Si3. ***C***. The holding current was linear with respect to the holding potentials used in this voltage-clamp experiment.

**Figure 9.**
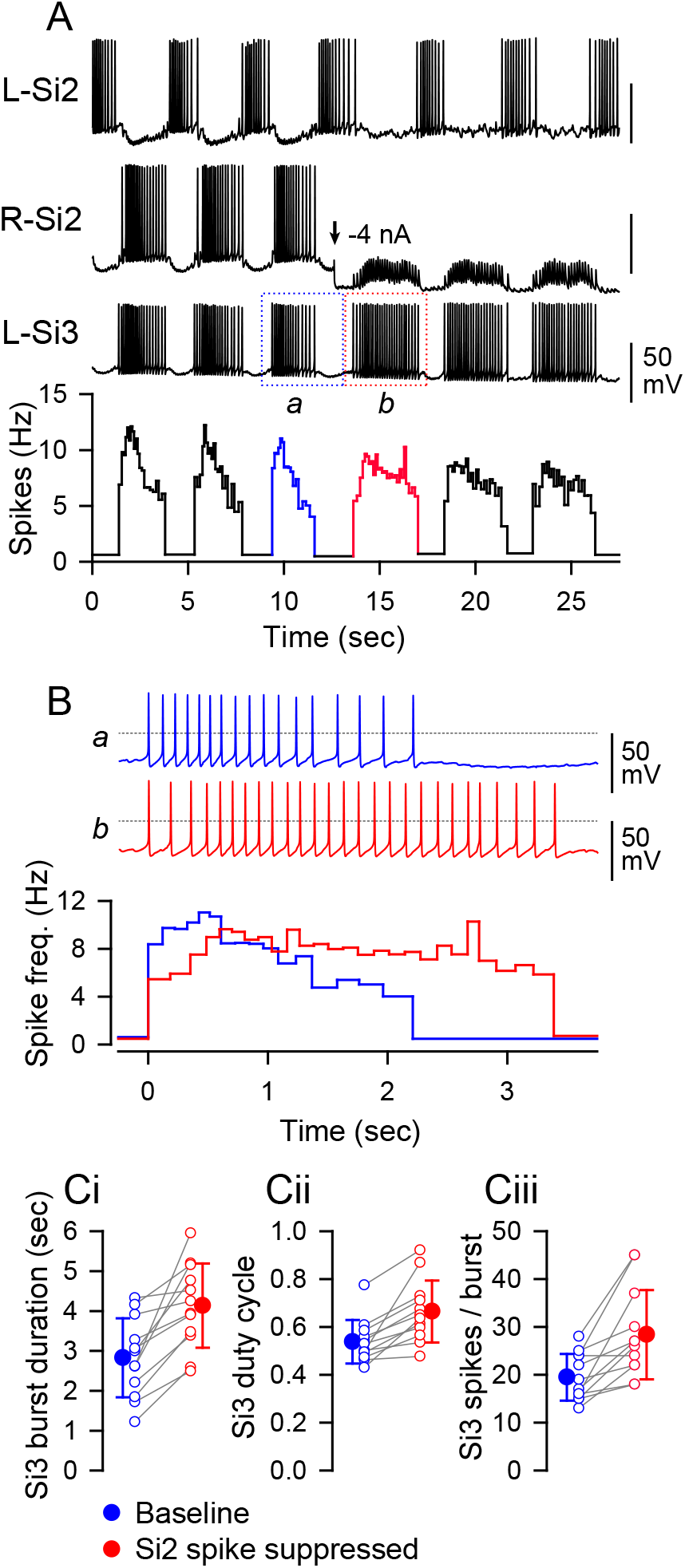
Suppression of the R-Si2 spiking prolonged Si3 bursts. ***A***. Simultaneous intracellular recordings of an Si2 pair and an L-Si3. A plot of instantaneous spike frequency is shown at the bottom. From the time indicated by the arrow, a hyperpolarizing current of −4 nA was applied to suppress the action potential firing of R-Si2. During the hyperpolarization, trains of EPSPs evoked by the L-Si3 spikes are visible in R-Si2. The bursts before and during the hyperpolarizing current injection are boxed by blue (a) and red (b) dotted lines. The spike frequency during the period is indicated by the same color in the plot below. *B*. A comparison of bursts before (***a*** – blue) and during the hyperpolarizing current injection (***b*** –red), expanded from ***A***. The dotted lines indicate 0 mV. ***C***. Comparisons of burst duration (Ci), burst duty cycle (Cii), and the number of Si3 spikes per burst (Ciii) before (blue) and immediately after (red) the suppression of the Si2 spikes by hyperpolarizing current injection. All three factors increased when the Si2 spikes were suppressed. Mean±SD represented by filled circle and error bars.

**Figure 10.**
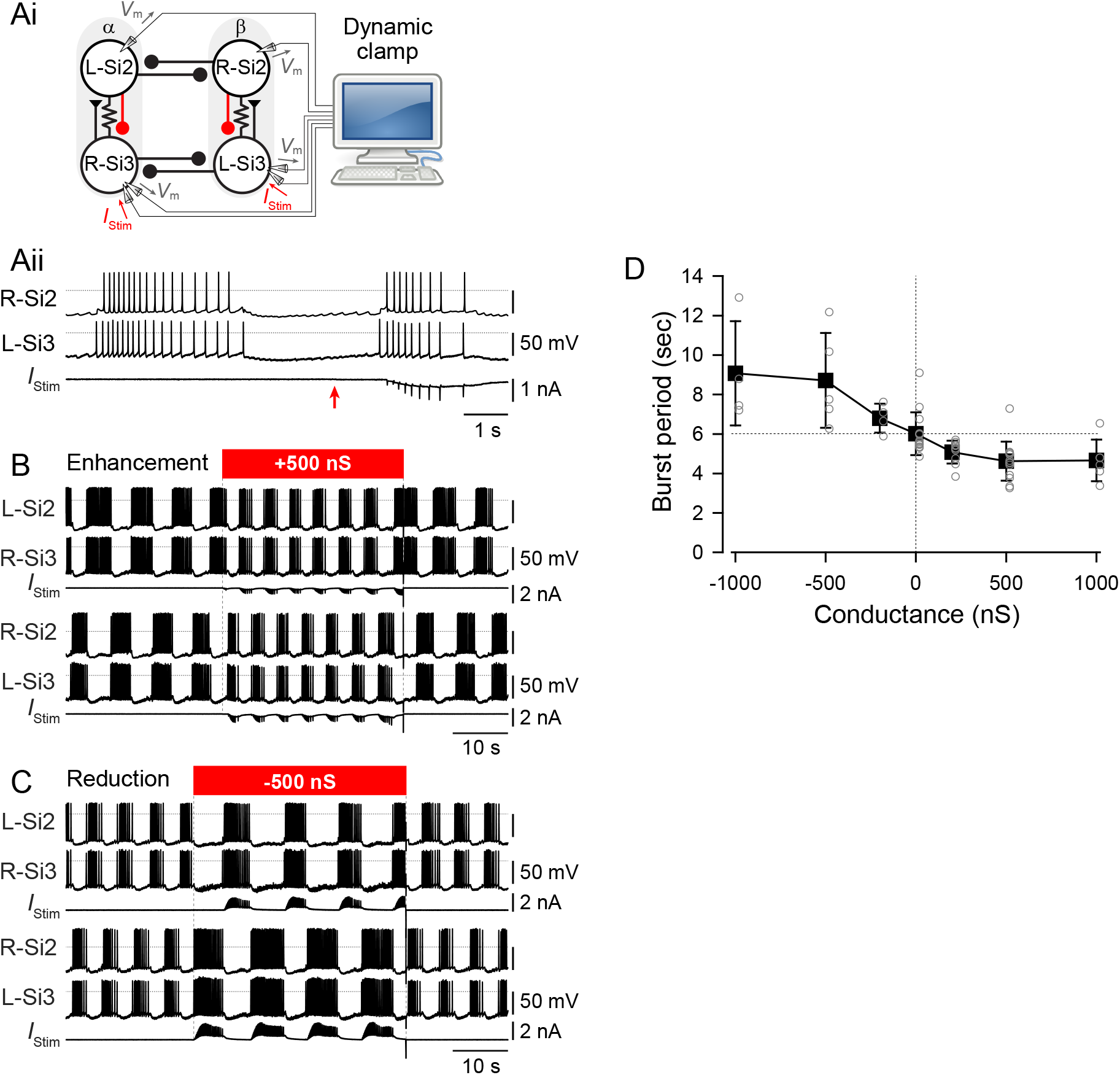
Artificial enhancement or suppression of slow inhibitory synaptic action of Si2 onto Si3 within each half-center module changed the burst period during a swim motor pattern. ***Ai***. The experimental arrangement for dynamic clamp. Every time the membrane potential (V_m_) of Si2 surpassed a specified threshold (50% of spike height), an artificial synaptic current (*I*_stim_) was calculated by the computer and injected into the contralateral Si3 to boost the existing Si2-to-Si3 slow synapse (red). Traces in ***Aii*** show the change in the burst before and during the application of dynamic clamp. The red arrow indicates the time of onset of dynamic clamping. The red arrow indicates the time when the dynamic clamp was started. The horizontal dotted lines show 0 mV. ***B***. Simultaneous intracellular recording from all four neurons shows the effect of dynamic clamping (red bar). The synaptic boost caused by adding artificial synaptic conductance (500 nS) with the dynamic clamp shortening the duration of bursts and decreased the period. ***C***. Synaptic suppression by counteracting IPSPs with a negative postsynaptic conductance to Si3 extended the burst duration and consequently increased the period. ***D***. Plot of burst periods against the conductance of artificial synapses from Si2 to Si3 in the same half-center module. Increasing the synaptic conductance reduced the burst period, whereas counteracting it extended the burst period.

The instantaneous activation, *S*(*t*) is given by the differential equation:

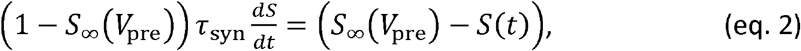

where

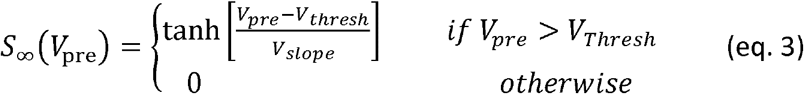

*S*_∞_ is the steady state synaptic activation and *τ*_syn_ is the time constant for synaptic decay. The time constant of the falling phase of the Si2-evoked slow IPSC in Si3 was 0.94 ± 0.2 sec (N = 5). In this study, *τ*_syn_ was set to 1.0 sec. *V*_pre_ is the presynaptic membrane potential of Si3 and *V*_thresh_ is the threshold potential for the release of neurotransmitter; it was set to the level of 50% height of the smallest Si3 action potentials. The synaptic slope parameter of the activation curve (*V*_slope_) was set to 25 mV. *g*_syn_ was varied between −1000 and 1000 nS.

### Statistics

Statistical comparisons were performed using SigmaPlot ver. 12.5 (Jandel Scientific, San Rafael, CA) for Student’s *t*-test, paired *t*-test, and Kruskal-Wallis one-way ANOVA on Ranks with all pairwise multiple Comparison Procedures. Shapiro-Wilk test was used to assume normality of data structure. In all cases, P<0.05 was considered significant. For *t*-tests, two-tailed P value was used. Results are expressed as the mean ± standard deviation (SD).

## Results

### Si2-evoked IPSPs had a more hyperpolarized reversal potential than those of Si3

In the *Dendronotus* swim CPG, two neurons, Si2 and Si3, each form inhibitory reciprocal synapses with their contralateral counterparts (Fig. 1B). When compared in high divalent cation (Hi-Di) saline, which raises spike threshold, thereby minimizing spontaneous and polysynaptic activity, the reversal potentials of the two synapses differed (Fig. 2). The average reversal potential of Si2-to-SI2 IPSPs was −73.9 ± 8.1 mV (N = 5; Fig. 2A*i*, B), whereas the Si3-to-Si3 IPSP reversal potential was −49.6 ± 6.5 mV (N = 6; Fig. 2A*ii*, B). There was a significant difference between these values (P < 0.001 by two-tailed t-test), suggesting that the currents underlying the IPSPs differ in ionic carriers. Thus, Si2 hyperpolarized its contralateral counterpart to a much greater extent than Si3 did when the postsynaptic neurons were near their resting potentials (about −50 mV).

### Si2 evoked a larger inhibitory effect on the opposite module than did Si3

During each cycle of the swim motor pattern, both Si2 and Si3 receive prolonged synaptic inhibition from their contralateral counterparts. To better evaluate the contributions of Si2 and Si3, we mimicked the activity during the swim motor pattern by stimulating each neuron independently with a 4 second train of action potentials (Fig. 3). When Si2 was stimulated, it caused a brief hyperpolarization and a delayed depolarization in both the Si2 and Si3 of the opposite module, which led to a post-inhibitory rebound (PIR) burst of action potentials in Si3 (Fig 3A). In 68% of those preparations (N=19 of 28), the contralateral Si2 also showed a PIR discharge, however, these were all triggered by the Si3-evoked EPSPs (Fig. 3A, asterisks). Thus, the inhibitory synaptic inputs from Si2 not only provided inhibition to the other half-center module, but they also induced an excitatory response in Si3 *via* PIR excitation.

In contrast, when Si3 was stimulated to produce a similar train of action potentials, as expected by the less negative reversal potential, the inhibitory synaptic potentials generated in Si2 and Si3 of the opposite module were markedly smaller than those generated by Si2 (Fig. 3B). Rebound firing in the contralateral Si3 occurred in only 69% of preparations (N=11 of 16), which induced the ipsilateral Si2 firing in 43.8% of those preparations (N = 7 of 16). Thus, Si3 was less effective than Si2 in causing a rebound discharge of Si3 in the opposite module.

The difference in the size of the evoked inhibitory potentials in response to a train of action potentials was more pronounced in Hi-Di saline (Fig. 4). A 4 sec, 10Hz spike train in Si2 hyperpolarized the contralateral Si2 by −11.7 ± 1.3 mV (N = 15, Fig. 4A,C). Si2 also hyperpolarized the ipsilateral Si3 by −6.4 ± 1.5 mV (N=15, Fig. 4A,C), because it is electrically coupled to the right Si2 (Sakurai and Katz 2016). In contrast, a 4 sec, 10 Hz spike train in Si3 hyperpolarized the contralateral Si3 by −2.2 ± 0.9 mV (N = 7) and the ipsilateral Si2 by −2.4 ± 1.0 mV (N = 9) (Fig. 4B,C). Thus, Si2 was more effective at inhibiting the other module than Si3 was.

### Si3, but not Si2, exhibited post-inhibitory rebound

After cessation of an Si2 spike train, Si3 in the other module not only recovered more quickly but also showed a PIR depolarization that exceeded above the original membrane, which was absent in Si2 (Fig. 4A, asterisk). Neither neuron exhibited rebound depolarization when Si3 was stimulated (Fig. 4B). The rebound depolarization induced by Si2 onto the Si3 in the other module was significantly larger than any other synaptic combination (Fig. 4D, P<0.05 by Kruskal-Wallis one-way ANOVA on Ranks with all pairwise multiple Comparison Procedures).

Si3 exhibited a larger PIR depolarization than Si2 following a hyperpolarizing current step (Fig. 5). In normal saline, cessation of a hyperpolarizing current pulse (4 sec, 1-4 nA) injected into Si2 produced few if any spike discharges (Fig. 5A*i*, B). In contrast, Si3 always exhibited a PIR discharge upon termination of the pulse (Fig. 5A*ii*, B). The number of spikes after the current pulse was significantly larger in Si3 than in Si2 (Fig. 5B; P < 0.001 by Two-way repeated-measures ANOVA with Holm-Sidak method, N = 9-12 for Si2 and N = 8-10 for Si3). With spike threshold elevated in Hi-Di saline, the membrane potential of Si2 quickly returned to resting potential when released from the hyperpolarizing current injection (Fig. 5C*i*), whereas Si3 showed a depolarizing overshoot (Fig. 5C*ii*, asterisk) that was significantly larger than seen Si2 (Fig. 5D; P < 0.001 by Two-way repeated-measures ANOVA with Holm-Sidak method, N = 20-23 for Si2 and N = 19-21 for Si3).

### Si3, but not Si2, exhibited a hyperpolarization-activated inward current

Si3 exhibited a larger “sag” potential than Si2 in which the membrane potential gradually depolarized during sustained injection of hyperpolarizing currents greater than −3nA (Fig. 5A, C, E; P < 0.001 by two-way repeated-measures ANOVA with Holm-Sidak method, N = 23 for Si2 and N = 21 for Si3). Under voltage clamp, Si3, but not Si2 exhibited a hyperpolarization-activated inward current (Fig. 6). Hyperpolarizing command voltage steps did not produce noticeable current response in Si2, whereas in Si3 they evoked slow inward currents at potentials more negative than −70mV (Fig. 6). Thus, Si2 and Si3 had distinct membrane properties; the hyperpolarization-activated slow inward current in Si3 likely underlies the sag depolarization and subsequent rebound depolarization, which produces the PIR discharge in Si3 and consequently induces the phase transition from one module of the half center to the other.

### Si2 evoked a slow synaptic inhibitory potential in Si3 within each module

We previously demonstrated that Si3 excites the Si2 in the same module through both a strong chemical synapse and strong electrical coupling, causing them to fire in near synchrony (Sakurai and Katz 2016). Here we found that Si2 inhibited Si3, causing it to stop firing action potentials when stimulated with a train (Fig 7A).

When spike threshold was increased in Hi-Di saline, an Si2 spike train produced a one-for-one electrotonic EPSPs in the contralateral Si3 (Fig. 7B*i*). With increased spike frequency, a slowly increasing hyperpolarizing potential counteracted the electrotonic EPSPs and produced an overall hyperpolarization (Fig. 7B*i*, 10 Hz). Application of Ca^2+^-free saline blocked this hyperpolarization without changing the electrotonic EPSPs (Fig. 7B*ii*, C). Subtracting traces in Ca^2+^-free saline (Fig. 7B*ii*) from those in Hi-Di saline (Fig. 7B*i*) revealed the Ca^2+^-sensitive component of the Si2-evoked postsynaptic potential in Si3 (Fig. 7B*iii*). Thus, Si2 was electrically coupled to Si3 and inhibited it through a slow potential caused by neurotransmitter release.

When Si3 was voltage-clamped, a spike train in the contralateral Si2 produced a slowly rising inhibitory postsynaptic current (IPSC) whose amplitude was outwardly rectifying (Fig. 8A,B). The reversal potential was variable across individuals ranging from −60 to −80 mV (Fig. 8B). The time constant of the falling phase of the slow IPSC was 0.94 ± 0.2 sec (N = 5). During the Si2 spike train, the membrane conductance of Si3 increased in all three preparations examined from an average of 80.7 ± 15.3 nS to 141 ± 22.1 nS (P = 0.039 by paired *t*-test, N = 3).

The outward rectification of the slow IPSC was not likely due to a change in background membrane conductance via opening or closing of voltage-gated channels, because in the absence of Si2 stimulation, the current/voltage relation was linear throughout the voltage range examined (Fig. 8C). At present, it is unclear how such apparent outward rectification was brought about; the variability in the reversal potential and the non-linearity of the I-V relationships could also be an experimental artifact caused by poor space clamping, suggesting that the site of synaptic input from Si2 is located contralateral to the Si3 soma.

### Feedback inhibition from Si2 to Si3 contributed to burst termination

To determine the role that the inhibition from Si2 to the contralateral Si3 plays in rhythmic bursting, we examined the effect of suppressing spikes in Si2 during the swim motor pattern (Fig. 9). When unperturbed, the intra-burst spike frequency of Si3 showed an initial peak and a subsequent steep decline during each burst (Fig. 9A,B, burst *a* - blue). Suppression of spiking in the co-active Si2 by injection of hyperpolarizing current immediately increased the duration of Si3 bursts (Fig. 9A,B burst *b* - red). The mean duration of Si3 bursts during the swim motor pattern increased significantly from 2.7 ± 0.9 sec to 4.1 ± 1.1 sec when the Si2 in its module was hyperpolarized to prevent it from spiking (Fig. 9C*i*; P < 0.001 by two-tailed paired t-test, N=12). Suppressing Si2 firing significantly increased both the mean duty cycle of Si3 bursts from 0.54 ± 0.09 to 0.67 ± 0.13 (Fig. 9C*ii*; P < 0.001 by Wilcoxon Signed Rank Test, N = 12) and the mean number of Si3 spikes from 19.5 ± 4.9 to 28.4 ± 9.3 (P < 0.001 by two-tailed paired t-test, N = 12) (Fig. 9C*iii*). Thus, although Si2 is electrically coupled to the Si3 in its module, the main effect of Si2 was to slowly decrease Si3 firing through a chemical inhibitory synapse.

### The magnitude of the slow inhibition from Si2 to Si3 played a role in setting burst period

The results suggest that the slow inhibition from Si2 to Si3 in each module decreases Si3 firing and hence deceases the excitatory drive from Si3 to Si2. We hypothesized that the self-inhibition within the module helps to terminate the burst and transition to activation of the other module. To test this, we systematically enhanced or reduced the magnitude of the inhibition from Si2 to Si3 in both modules during a swim motor pattern using the dynamic clamp technique. The amplitudes and time courses of the Si2-evoked synaptic currents were calculated in real time (see Materials and Methods) and injected into Si3 through the recording electrode in response to spikes in the contralateral Si2 (Fig. 10A*i*). While the dynamic clamp was engaged, an Si2 burst set off a slow negative current injected into Si3, boosting the slow inhibitory synapse, which shortened the Si3 burst (Fig. 10A*ii*). Spikes in the postsynaptic Si3 also produced spike-like downward surges in the injected current, because the spikes moved the membrane potential away from the reversal potential of the slow IPSC (−70 mV), increasing the driving force for the current.

Augmenting the amplitude of the Si2-to-Si3 slow inhibitory synaptic conductance shortened the Si3 burst duration, thereby decreasing the burst period (Fig. 10B,D). In contrast, counteracting the inhibition with an inverted conductance reduced the Si2-to-Si3 inhibition and extended Si3 bursts, thereby increasing the burst period (Fig. 10C,D). Thus, the feedback inhibition from Si2 to Si3 within each module plays a crucial role in maintaining the periodicity of the half-center oscillator.

## Discussion

The results presented show that stable alternating bursting in the *Dendronotus* swim CPG is an emergent property of the network that arises from the synaptic actions and the membrane properties of the four neurons, with the two neurons in each module playing complementary roles in burst initiation and termination (Fig 11). Specifically, we found that synaptic inhibition of Si2 by its contralateral counterpart in the opposite module provides the hyperpolarization needed for the electrically-coupled Si3 to rebound and fire action potentials. The spikes in Si3 then excite Si2 causing both neurons to fire a maintained train of spikes. Si2 spiking slowly inhibits Si3, eventually terminating the burst and releasing the opposite module for the next burst. This creates a sequence of activation and feedback inhibition in which each of the four neurons in turn becomes dominant (Fig. 11 B,C).

**Figure 11.**
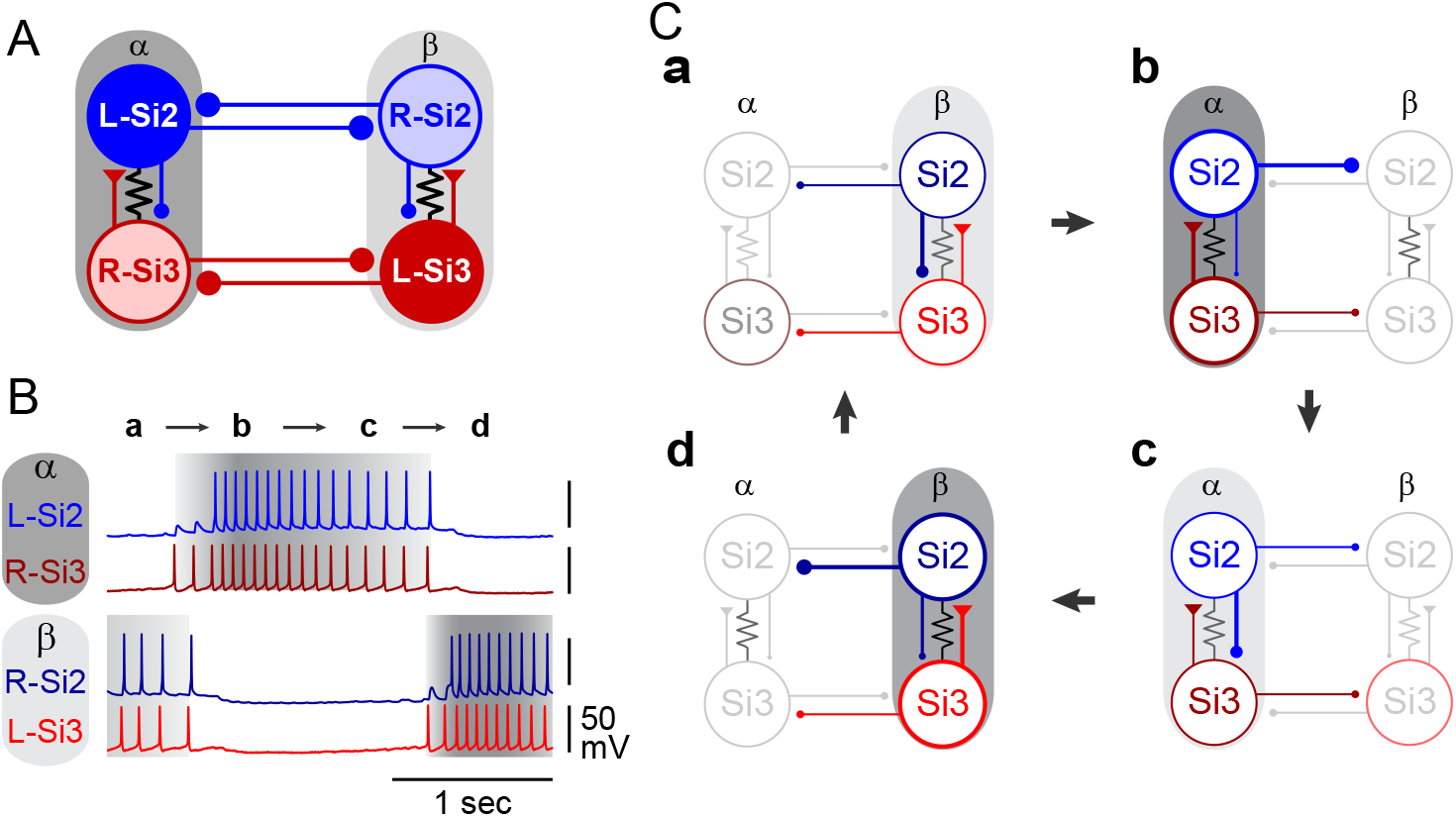
Sequence of activities of the CPG neurons. ***A***. The updated connectivity of the *Dendronotus* swim CPG. In each module (α and β) Si3 makes an excitatory synapse on the contralateral Si2, which returns a slow inhibitory synapse onto the Si3. Lines terminating in triangles indicate excitatory synapses, filled circles inhibitory synapses. Resistor symbols indicate electrical connections. ***B***. The activity of a circuit neuron is divided into four parts (**a, b, c, d**) shown schematically in **C.** In **Phase a,** Si2 inhibition of Si3 increases, causing the firing frequency of Si2 and Si3 in module β to decline, thereby decreasing inhibition of module α. **Phase b,** Si3 in module α starts firing, and Si2, which is driven by Si3, begins to fire a little later. **Phase c.** Si2 inhibition of Si3 increases, causing the firing frequency of Si2 and Si3 in module α to decline, thereby decreasing inhibition of module β. **Phase d.** Si3 in module β escapes from inhibition and starts firing together with Si2 and suppressing activity in module α.

### Feedback inhibition as a mechanism of release in a HCO

A key finding of this study is the discovery of a synaptic mechanism that serves as a brake on self-excitation within each module of the half-center circuit. In a previous study, we found that Si3 excites the Si2 in its module with very large EPSPs and the two neurons are tightly coupled by electrical synapses, causing synchronized activity that is essential for maintaining half-center oscillator activity (Sakurai and Katz 2016). Here, we found that Si2 evokes slow inhibitory potentials in Si3 that summate until Si3 stops firing, thereby ending the burst. The magnitude of this feedback inhibition sets the burst duration of the HCO as shown with dynamic clamp manipulation.

Previously, Friesen (1994) suggested that spike frequency adaptation of each half-center module or fatigue of mutually inhibitory synapses are the potential mechanisms that release the other module from inhibition. However, few such circuits have been reported; instead, neural circuits with a half-center configuration often are capable of generating endogenous bursting upon isolation (see however: Arshavsky et al. 1986; Cymbalyuk et al. 2002; Selverston and Miller 1980). To our knowledge there have been no prior reports of inhibition within a module playing an essential role in other HCOs. The division of labor between Si2 and Si3 within each module allows the summation of synaptic inhibition from Si2 to Si3 to serve as a mechanism that determines burst duration.

### Distinct roles for contralateral inhibition

Although both Si2 and Si3 had reciprocal inhibitory synapses with their contralateral counterparts, as is typical for a half-center oscillator, only the Si2 synapses were responsible for triggering PIR in the opposite module. The Si2-Si2 IPSPs were larger than the Si3-Si3 IPSPs because they had a more negative reversal potential. The Si2 spike train hyperpolarized Si3 in the opposite module, but this action was indirect through electrical coupling from Si2 to Si3 within that module. The inhibitory effect of the Si3-evoked IPSPs was mostly due to shunting inhibition caused by an increase in membrane conductance because their reversal potential was close to the resting membrane potential (Paulus and Rothwell 2016).

The synaptic actions of Si2 and Si3 differed depending on the postsynaptic cell, suggesting that the postsynaptic neurons differ in receptor type. The Si2-Si2 synaptic potentials were rapid, with a reversal potential suggestive of a ligand-gated potassium channel. The Si2 to Si3 synaptic potentials also had a reversal potential near potassium equilibrium, but the fact that they did not appear as discrete synaptic potentials corresponding to the presynaptic spikes and their slow kinetics suggests that they are mediated by a G-protein coupled receptor. The Si3-Si3 synaptic properties are consistent with being mediated by a ligand-gated chloride channel. The Si3-Si2 synapse appears to be mediated by a ligand gated cation channel with a reversal potential estimated to be near 0 mV (Sakurai and Katz 2016). Both effects of Si3 are blocked by curare (Sakurai and Katz 2017), suggesting that they are mediated by different variant of nicotinic acetylcholine receptors, which can be cation or anion selective in molluscs (Jiao et al. 2019; van Nierop et al. 2006; van Nierop et al. 2005).

### Different PIR properties for burst initiation

Only the Si3 each module showed PIR when released from hyperpolarization or after a train of IPSPs evoked by the other half-center module. The mechanism for the PIR was a hyperpolarization-activated inward current, which was absent in Si2. This difference in membrane properties ensures that Si3 fires before Si2 during each burst. This may contribute to the stability of bursting by inserting a delay before feedback inhibition from Si2 can occur.

The sag potential that develops as a result of accumulation of the hyperpolarization-activated inward current may contribute to the escape of these neurons from synaptic inhibition from the opposing module as seen in other systems. (Arbas and Calabrese 1987; Pirtle et al. 2010). Thus, the *Dendronotus* HCO has elements of both release and escape mechanisms as suggested by the occasional overlap of the first Si3 spike and the last Si2 spike of the opposite module (for example, Fig. 11B)

Active membrane properties of neurons, such as “sag” depolarization and PIR, have been shown to be involved in the transition of activity by allowing the postsynaptic neurons to escape from inhibition or firing once released from inhibition (Arbas and Calabrese 1987; Arshavsky et al. 1985; Dethier et al. 2015; Li and Moult 2012). Studies using computer models also demonstrated the role of PIR in escaping from inhibition (Daun et al. 2009; Perkel and Mulloney 1974; Wang and Rinzel 1992). Here we found that one neuron in each module had this function.

### The Dendronotus swim CPG is a true network oscillator

Many neural circuits with a half-center configuration are capable of generating endogenous bursts upon isolation of the two modules (Arshavsky et al. 1986; Cymbalyuk et al. 2002; Selverston and Miller 1980). Similarly, conditional pacemaker properties also have been found in each half of the locomotor circuit in the spinal cord (Kiehn et al. 1996; Li et al. 2010; Reith and Sillar 1998; Wallen and Grillner 1987). When the left and right modules of the spinal swimming circuit are separated in lamprey, each module exhibits rhythmic bursting activity at a faster rhythm than normal (Grillner 2003). If each half-center possesses endogenous rhythmogenesis, then, the role of mutual inhibition between half-centers is limited to ensuring coordinated bursts occur in alternation (Ausborn et al. 2018; Hagglund et al. 2013). A counterexample to this was reported by Moult et al. (2013), in which they showed that the onset of rhythmic activity did not occur instantaneously when unilateral activity is momentarily inhibited, but took some time, suggesting that some neuroplasticity may be involved in the experimentally induced onset of unilateral automaticity.

In *Dendronotus*, it is unlikely that each module of the swim CPG has rhythmogenesis because no oscillatory activity was seen when the activity of one side was suppressed by current injection (Sakurai and Katz 2016). Furthermore, blocking the synaptic actions Si3 with curare halts all rhythmic activity (Sakurai and Katz 2017). The Si3 pair alone cannot generate oscillatory activity even when depolarized by current injection. Thus, the *Dendronotus* swim CPG is a unique example of a network oscillator with a half-center configuration, in which none of the neurons, nor the opposite modules are capable of rhythmogenesis; rhythmic activity is an emergent property of the network as a whole.

### All synaptic connectivity in the pedal ganglia is contralateral

It is worth noting that almost all of the synapses made by pedal ganglion neurons in both *Dendronotus* all other nudibranchs studied are between contralateral neurons; very few synapses have ever been found between pedal ganglion neurons whose somata are ipsilateral to each other (Sakurai et al. 2014a; Sakurai and Katz 2016). In the nudibranch, *Melibe leonine*, there is an additional pedal swim neuron (Si4), which also has only contralateral synapses (Sakurai et al. 2014a). We are aware of only one connection between ipsilateral pedal ganglion neurons in any nudibranch species studied; ipsilateral Si2 was shown to elicit one-for-one EPSPs in pedal efferent neurons in *Melibe* (Thompson and Watson 2005). This in contrast with cerebral ganglion neurons, such as Si1 the DSIs and C2, which make both ipsilateral and contralateral connections (Gunaratne et al. 2017; Katz 2009; Sakurai et al. 2014a; Sakurai and Katz 2019).

The lack of synaptic connectivity within the pedal ganglion may extend to all gastropods studied. There are reports of “non-synaptic”, volume transmission in the pteropod *Clione limacina* (Arshavsky et al. 1988) and a slow inhibitory synapse between two pedal neurons in *Lymnaea stagnalis* (Inoue et al. 1996), however there are no reports of one-for-one fast synaptic potentials between neurons within the pedal ganglion for any gastropod. This suggests something fundamental about the organization of gastropod nervous systems, namely that aside from the limited number of contralaterally-projecting interneurons, the majority of pedal ganglion neurons are efferent, projecting to the rest of the body.

Thus, the organization of the two modules being composed of contralateral Si2 and Si3 may be a result of a constraint on the system. The CPG could not be composed neurons on the left acting together because they do not synapse on each other. Thus, the only way to construct a half-center oscillator is to have each half composed of contralateral neurons.

### Comparisons to the Melibe swim CPG

The nudibranch, *Melibe*, swims with rhythmic left-right body flexions like *Dendronotus*. The swimming behaviors are likely homologous because all species within the clade swim in this manner (Sakurai and Katz 2017). The *Melibe* swim CPG, which consists of 8 neurons (left and right) Si1, Si2, Si3, and Si4, is also organized into a half-center configuration, but the roles of Si2 and Si3 differ from those in *Dendronotus* (Sakurai et al. 2014a; Sakurai and Katz 2017; 2016; Sakurai et al. 2014b). The primary half-center kernel, consists of Si1, Si2, and Si4, which can generate alternating burst discharges without Si3. The role of Si3 is to terminate the bursts of each module.

Thus, homologous neurons have opposite functions in the two HCOs; in *Dendronotus*, Si3 initiates bursts and Si2 terminates them, but in *Melibe* S2 is part of the burst generator and Si3 terminates the bursts. These differences arise in part due to differences in the synaptic actions and connectivity of the homologous neurons. In addition, Si1, which is part of the HCO in *Melibe*, plays an extrinsic neuromodulatory role in *Dendronotus*, enhancing the synaptic actions of Si3 (Sakurai and Katz 2019).

The direction of evolutionary change is not clear. We do not know if the more complex eight cell *Melibe* CPG arose from a simpler ancestral *Dendronotus*-like circuit or whether the *Dendronotus* swim CPG arose as refinement of a larger *Melibe*-like circuit. Regardless of how it arose, the *Dendronotus* swim CPG is beautiful in its simplicity; two neurons with complementary synaptic actions and membrane properties partition the functions needed to ensure stable alternating bursts.

## Acknowledgments

This work was supported by NSF grant IOS-1120950.

